# Impact of global change on future Ebola emergence and epidemic potential in Africa

**DOI:** 10.1101/206169

**Authors:** D. W. Redding, P. M. Atkinson, A. A. Cunningham, G. Lo Iacono, L. M. Moses, J. Wood, K. E. Jones

## Abstract

Animal-borne or zoonotic human diseases (e.g., SARS, Rabies) represent major health and economic burdens throughout the world, disproportionately impacting poor communities. In 2013-2016, an outbreak of the Ebola virus disease (EVD), a zoonotic disease spread from animal reservoirs caused by the Zaire Ebola virus (EBOV), infected approximately 30,000 people, causing considerable negative social and economic impacts in an unexpected geographical location(Sierra Leone, Guinea, and Liberia). It is not known whether the spatial distribution of this outbreak and unprecedented severity was precipitated by environmental changes and, if so, which areas might be at risk in the future. To better address the major health and economic impacts of zoonotic diseases we develop a system-dynamics approach to capture the impact of future climate, land use and human population change on Ebola (EVD). We create future risk maps for affected areas and predict between a 1.75-3.2 fold increase in EVD outbreaks per year by 2070. While the best case future scenarios we test saw a reduction in the likelihood of epidemics, other future scenarios with high human population growth and low rates of socioeconomic development saw a fourfold increase in the risk of epidemics occurring and almost 50% increase in the risk of catastrophic epidemics. As well as helping to target where health infrastructure might be further developed or vaccines best deployed, our modelling framework can be used to target global interventions and forecast risk for many other zoonotic diseases.

**Significance Statement:** Despite the severe health and economic impacts of outbreaks of diseases like SARS or Zika, there has been surprisingly little progress in predicting where and when human infectious disease outbreaks will occur next. By modelling the impacts of future climate, land use and human population change on one particular disease Ebola, we develop future risk maps for the affected areas and predict 1.7-3.2 times as many human Ebola outbreaks per year by 2070, and a 50% increase in the chance that these outbreaks will become epidemics. As well as helping to target where health infrastructure might be further developed or vaccines deployed, our approach can also be used to target actions and predict risk hotspots for many other infectious diseases.

## Introduction

Little is known about how the majority of human infectious diseases will be affected by predicted future global environmental changes (such as climate, land use, human societal and demographic change) (1–5). Importantly, two thirds of human infectious diseases are animal-borne (zoonotic) (6) and these diseases form a major, global health and economic burden, disproportionately impacting poor communities (7, 8). Many zoonotic diseases are poorly understood, and global health responses to them are chronically underfunded (9). The 2013–2016 Ebola outbreak was unprecedented in terms of size, financial cost, and geographical location (10, 11); a stark illustration of our knowledge gaps, and demonstrating that it is imperative we develop quantitative approaches to better forecast zoonotic disease risk.

Ebola virus disease (EVD) was first identified in 1976, and since then there have been approximately 23 recognized outbreaks (12), predominantly within central Africa. EVD is caused by any one of four pathogenic strains of Ebola virus:Zaire (EBOV), Sudan (SUDV), TaïForest (TAFV), and Bundibugyo (BDBV). It presents as a non-specific febrile illness that can cause haemorrhagic fever, often with a high case fatality rate in diagnosed patients (13).Some Old World fruit bat species (Family Pteropodidae) have been suggested as reservoir hosts (14), however, while there is limited direct evidence, they are strong candidates to playa key role either as an reservoir or amplifying host (15, 16). In areas with EVD, there are frequent direct and indirect human-bat interactions, e.g., via bush meat hunting and during fruit harvesting (17), presenting numerous opportunities for bat-to-human pathogen spill-overs to occur. Additionally, a third of known zoonotic spill-overs have been connected to contact with great apes and duikers, although there is no evidence that these species act as reservoir hosts (10). It is clear, however, that once spill-over occurs human social factors such as movement and healthcare responses greatly influence the cumulative outcome of an outbreak (18). For instance, previous work has highlighted the importance of family interactions (19), funeral practices (20) and differential transmission rates in hospitalized individuals (18).

Many attempts to understand Ebola outbreak dynamics have focused on mechanistic modelling approaches of human-to-human transmission post spill-over from animal hosts (13, 18, 19, 21–24). Mechanistic, or process-based, models are ideal for capturing epidemiological characteristics of diseases and, importantly, testing how disease outbreaks might be impacted by intervention efforts (25). One downside is that mechanistic models rarely incorporate spatially heterogeneous ecological and environmental information (26), such as the known high variance of bat abundance and pathogen sero-prevalence across widespread individuals (27). In this context, correlative, or pattern-based, models (e.g.MaxEnt, Boosted-regression trees) have been used to simultaneously capture the spatial risk of both zoonotic spill-over and subsequent human-to-human infection (12). For some spatially-explicit analyses, there have been attempts to incorporate spatial patterns of human populations (28), while other have included air transportation networks (29),but no studies that we are aware of have considered whole-system analyses for major epidemic zoonoses, such as Ebola. Like other rare or poorly-sampled diseases, Ebola suffers from limited data availability, meaning pattern-finding, correlative analytical techniques are at adisadvantage (30).

In 2014 a spill-over in Gueckedou district, Guinea of Ebola-Zaire virus led to an EVD outbreak approximately 100 times larger than any of the previous 21 known outbreaks (31). Such epidemics have a disproportionate impact on the affected societies. For example, the World Bank estimates a cost of US$2.2 billion to the three most affected countries (32) due to, amongst other drivers, widespread infrastructure breakdown, mass migration, crop abandonment and arisein endemic diseases due to overrun healthcare systems. Recent work has uncovered the human-to-human transmission patterns underlying this outbreak, using case (33) and genomic data (31) to demonstrate that EVD spread can be successfully predicted by a dispersal model that is weighted by both geographic distance and human population density. Attempting to understand zoonotic epidemic risk, however, using a human-only transmission model and without incorporating host ecology would inevitably lead to areas with high human density and connectivity being identified as the regions with the highest risk, despite some areas of these lacking competent hosts. Therefore, to model both the spatial variation in spill-over risk and, concurrently, the likely progression of subsequent outbreaks in human populations, we need to take a system-dynamics modelling approach (1, 34). Key non-linear feedbacks can also be captured, for example, the trade-off between increasing human populations and loss of reservoir host species through anthropogenic land-use conversion, and using this to design the optimal roll-out of vaccinations (35) and other interventions in a changing landscape.

Here, we use a disease system-dynamics approach (Figure 1) to extend a discrete-time, stochastic epidemiological compartmental model incorporating spatial environmental variability (Environmental-Mechanistic Model or EMM, Figure 2) to simulate present day spill-over and subsequent human-to-human transmission of the Zaire Ebola virus (EBOV) (the strain responsible for the 2013-2016 outbreak in West Africa). We model the impact of future anthropogenic changes on the occurrence and spread of this disease in 2070 (36–38) under a variety of possible integrated global change scenarios (39). We use a combination of three Representative Concentration Pathways (RCP) scenarios of increasing greenhouse gas concentrations: RCP4.5, RCP6, and RCP8.5 (40), and three possible socio-economic development scenarios (Shared Socio-economic Pathways or SSP), ordered by increasing human population density and reduced regional socio-economic cooperation:SSP1, SSP2 and SSP3. Finally, we compare the changes to spatial patterns of risk and chancesof outbreaks and epidemics occurring across Africa.

**Fig. 1.**
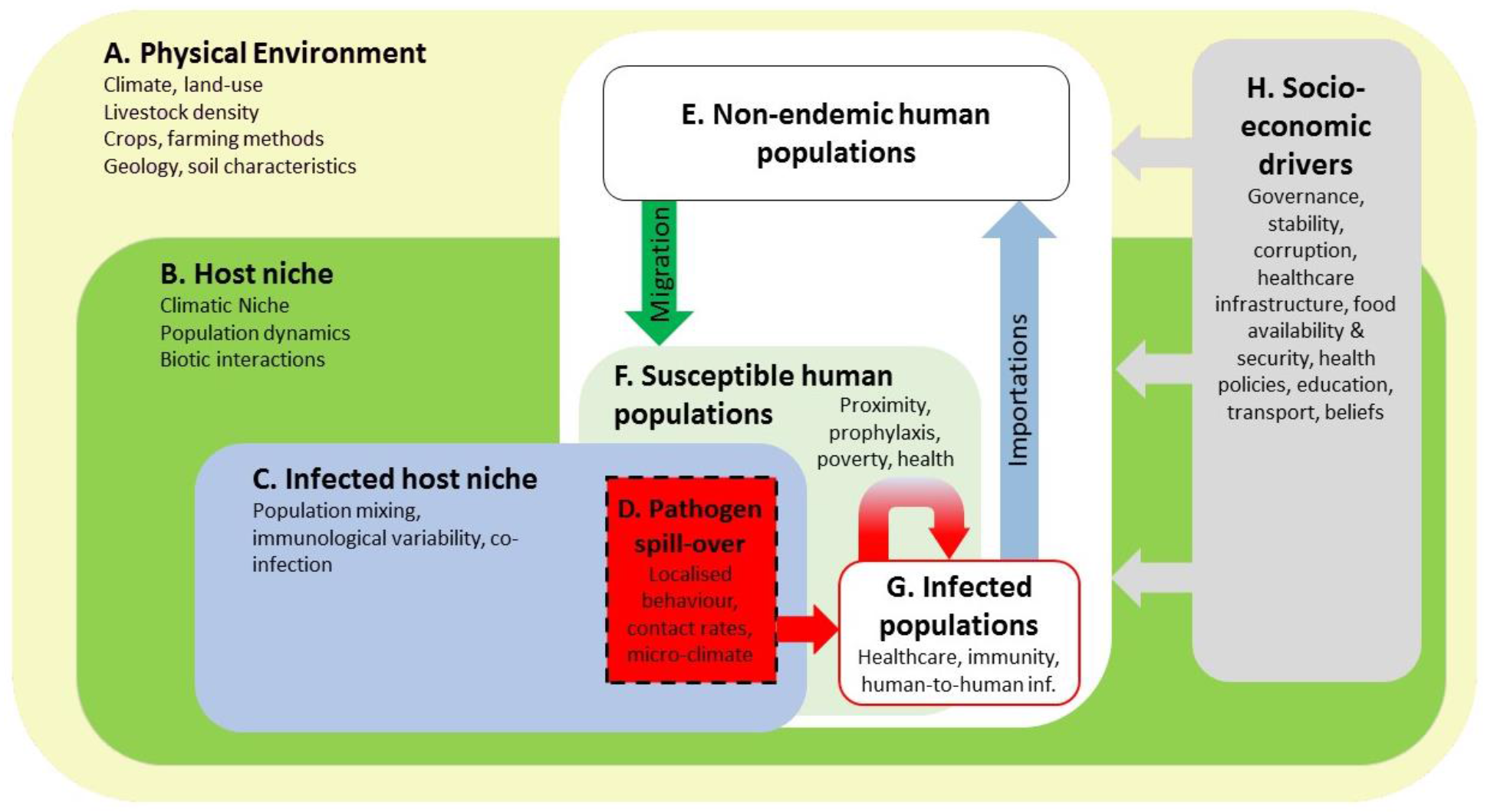
System-dynamics model of zoonotic disease transmission. Letters A-H indicate major system components, arrows showing links, and key sub-components in smaller font.

**Fig. 2.**
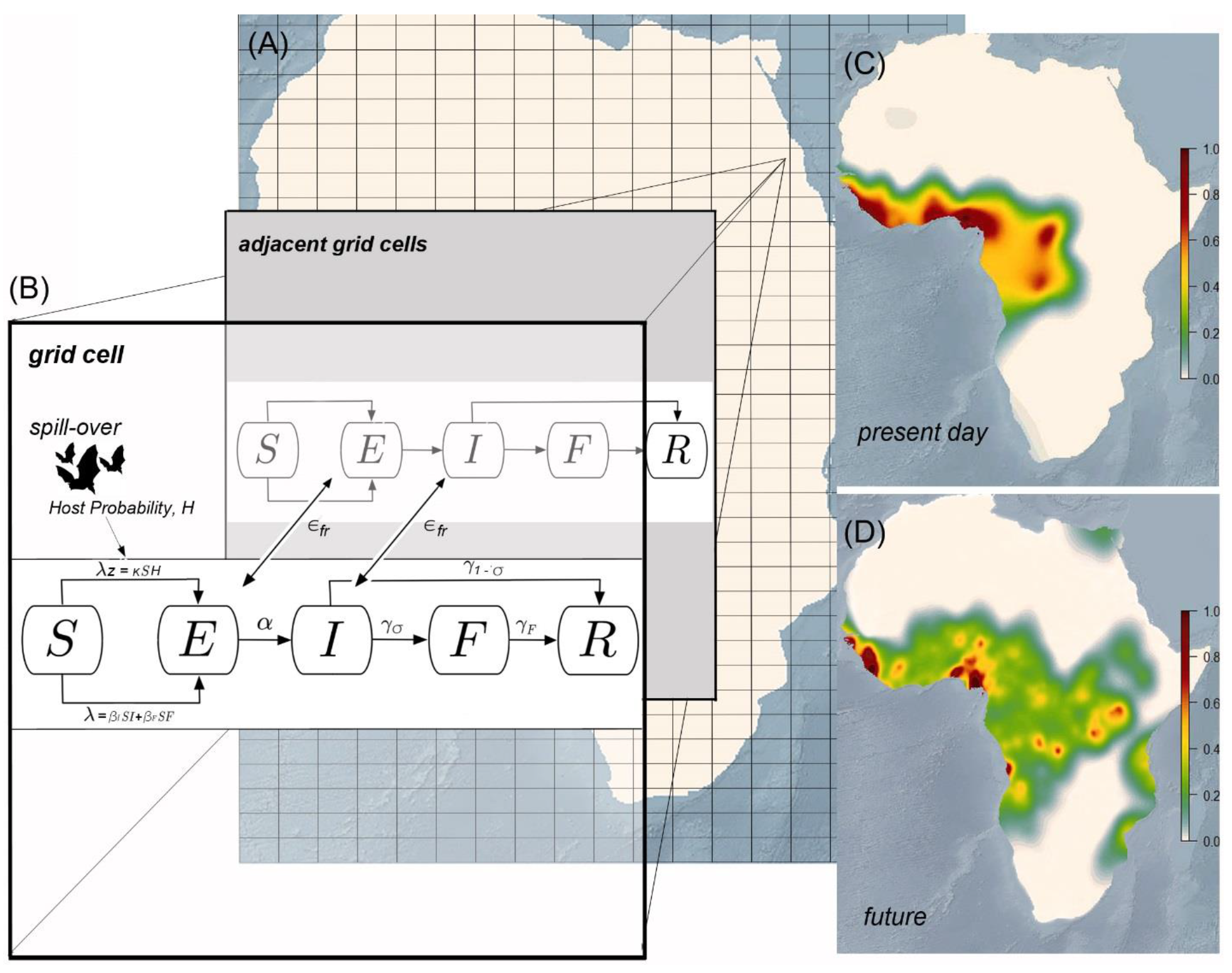
Predictive Integrated Zoonotic Model (EMM) EBOV Simulation Schematic. Within 0.0416° grid cells across the globe (**A**), we used a SEIFR (Susceptible, Exposed, Infectious, Funeral and Removed) disease compartmental model (**B**), to estimate the number of people in each compartment. S-E transmission rate was determined for each grid cell by calculating the force of zoonotic infection (between hosts and humans) *λ*_z_, and within human populations f (see Materials and Methods). Travel of exposed or infectious individuals between grid cells occurred across existing road and flight transport networks, with transmission rate *ε*_*fr*_. Mean transition rates used as the starting parameters for simulations wereas follows: *α* for E-I was calculated as the reciprocal of incubation time in days (*α*=1/7), *γ*_*σ*_ (I-F transition rate) was the product of the probability of the reciprocalofdays infectious(*γ*=1/9.6) and poverty-weighted case fatality rate (*σ*=0.78), *γ*_1−*σ*_ (I-Rtransition. rate) was the product of the probability of the reciprocal of days infectious (*γ*=1/9.6) and probability of recovering (1−σ), and *γ*_*ϝ*_(F-R transition rate) was the reciprocal of the burial time of 2 days. Each simulation was run 2500 times for 365 days in each grid cell containing a humanpopulation. The total number of people in each compartment per grid cell, per day from each simulation was then used to calculate the total number of index and secondary cases and mapped spatially (C). Bioclimatic, land use and demographic conditions werethen changed to predicted values for 2070 to estimate changes to *λ* and *λ*_*ᴢ*_, and the simulations repeated toinvestigate impacts of global change on disease (D).

## Results

Our EMM simulation for present day EBOV-EVD risk correctly identified areas of observed outbreaks as high risk, such as Democratic Republic of Congo, Gabon and the 2013-2016 outbreak in West Africa, but also identified some areas where EVD has not been reported, such as Nigeria and Ghana (Figure 3A). As a result, our model suggests that the at-risk area for EBOV-EVD is much larger than the areas known to have experienced disease outbreaks thus far. Our risk map also identified areas that are endemic for the other EVD strains,likely due to similar transmission pathways and reservoir host characteristics (Figure 3A). Although the index case risk map (Figure 3B) shows a similar spatial pattern to all cases, high risk spill-over areas are constrained to more distinct hot-spots. Importantly, the locations of index cases that resulted in epidemics were even more geographically constrained, with G ana, Sierra Leone, Liberia, Kenya Uganda and Cameroon all having medium risk but Nigeria is the focus of the highest potential for epidemic spill-over (Figure 3C). Comparing the mean number of spill-overs per year gave higher results for present day simulations with 2.464 spill-overs per year (95% CI 2.361-2.567) compared to the mean historical number over the last 40 years: 0.75 (95% CI 0.695-0.905). High risk of Ebola case importation using the current network of airline flights was seen in China, Russia, India,the United States as well as many high-income European countries (Figure 4). Especially high importation risk, however, was seen in Italy and Germany.

**Fig. 3.**
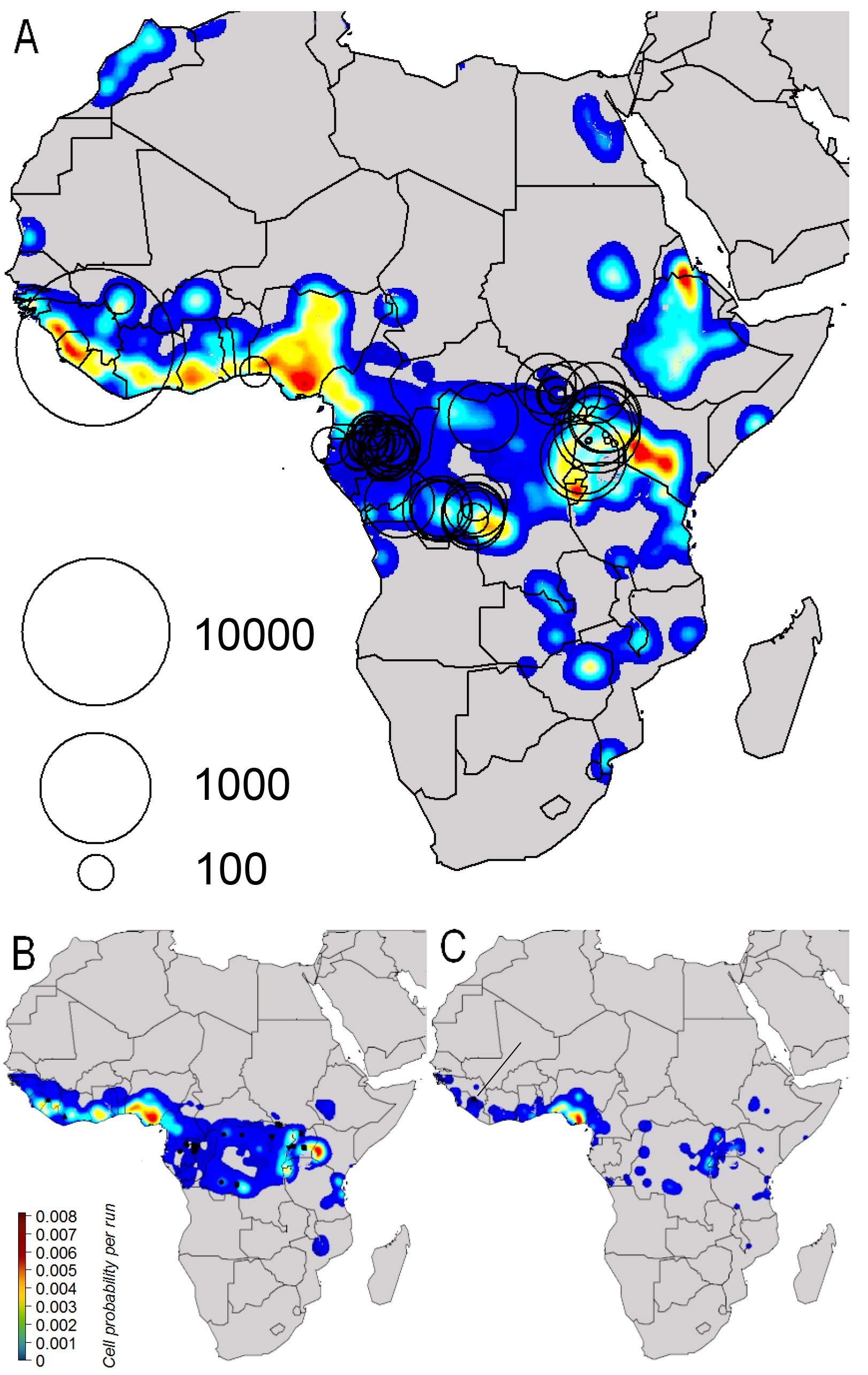
Present day risk for Zaire Ebola virus (EBOV) from EMM simulations. Maps represent the proportion of times between zero (dark blue) and 0.01 (red) when a EVD-EBOV case waspresent in a grid cell (0.0416a) across 2500, 365 day simulation runs for the presentday, where (A) shows all cases (both index and secondary), (B) index cases only, and (C)index casesfrom epidemics (1500+ cases). Black open circles in (A) represent logoutbreak size with the location of the index case at the centre of the circle. Black symbols in (B)representalllocations of known EVD index cases from different viral strains, wherecircles represent Zaire (EBOV), square Sudan (SUDV), triangles Tai Forest (TAFV), and tetrahedrons Bundibugyo (BDBV).Single black circle in (C) shows the only known site where an epidemic has occurred, with the black line highlighting its location

**Fig. 4.**
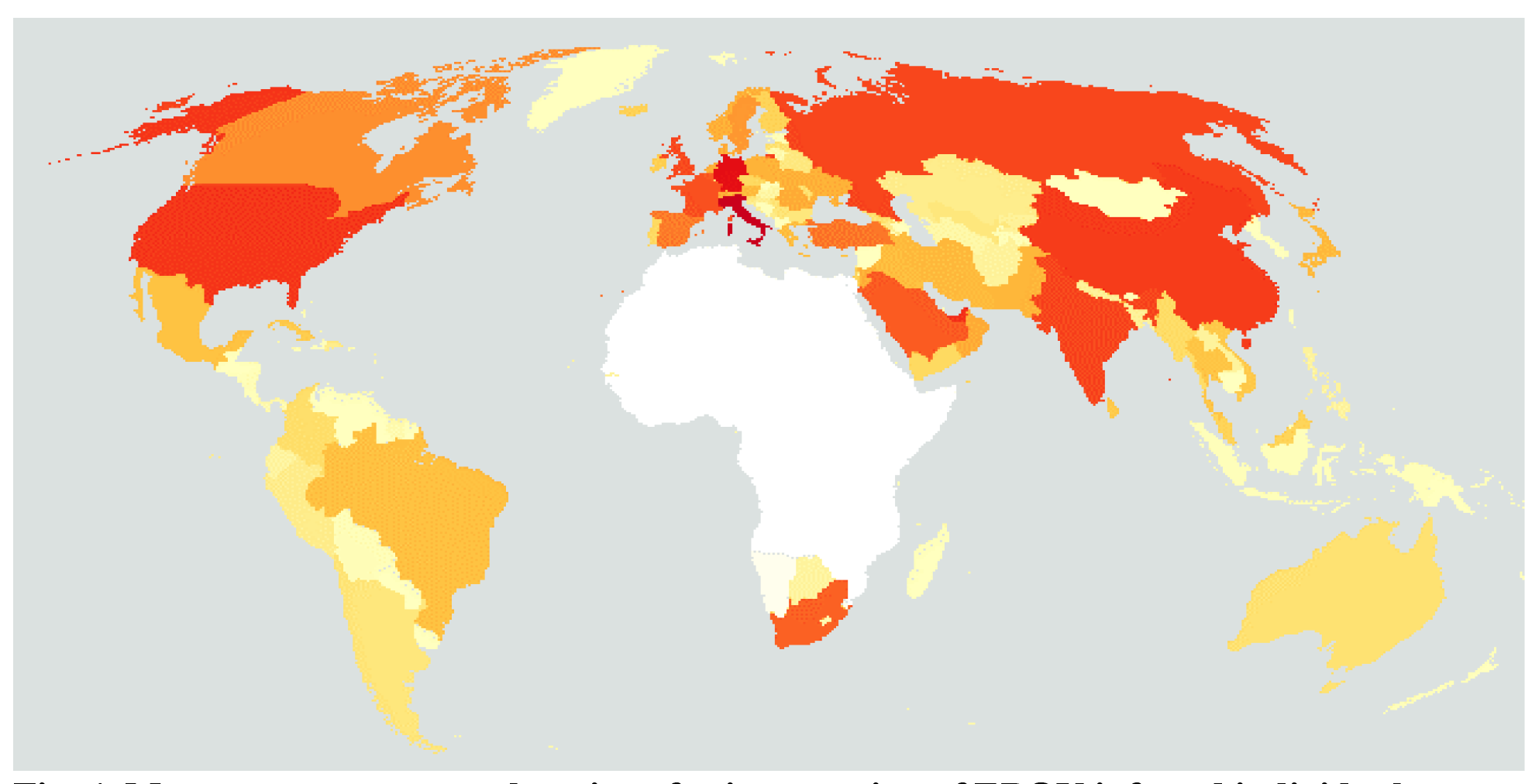
Most common country locations for importation of EBOV infected individuals. Map shows the countries that received, by airline flights, the most EBOV infected individuals (Red) with paler, orange and then yellow coloured countries having proportional fewer importations and white showing the EVD endemics area. Data come from 2500 simulations of EVD outbreaks under present data climate, land-use, demographic and transportation conditions.

Similar to historic data (Figure S3), the distribution of the final size of the simulated outbreaks was multimodal with distinct peaks at very low numbers (less than 3 cases) and medium outbreaks (approximately 3-1500 cases) (Figure S3).Through extensive simulations we were able to explore the lower probability areas of the distribution effectively and, unique to the simulation data, there is a third peak of outbreaks(here we term ‘epidemics’) with high, to very high, numbers of cases (1500-100,000,000 cases). This threshold of assigning an outbreak with greater than 1500 cases as an epidemic also corresponds to the top 1 percentile of a log-normal distribution approximating the variation in pre-2016 observed outbreak sizes (~1538 cases per year). Of the ~2500 simulation runs for present day conditions, epidemics (>1500) occurred approximately in 5.8% of the yearly simulations, with catastrophic epidemics (>2,000,000) occurring in around 2.3%of simulations, or once every 43.5 years given current conditions. From the sensitivity testing, the key parameters that affected outbreak size were illness length and R0, which positively increased case numbers (Figure S4a), where as the annual spill-over rate (Figure S4a) was most impacted by the spill-over rate constant (strongly positive), shape of the poverty/spill-over curve (weakly positive), and by host movement distance (weakly negative).

Our future EMM simulations estimate an annual increase in maximum area impacted by the disease from 3.45 million km^2^ to 3.8 million km^2^ under the scenario by 2070, with increases in maximum area seen under all future scenarios (Figure 5A, Figure 5D, Figure 5G). The maximum areas where spill-overs could occur, however, increased by just 1% under the RCP4.5 SSP1 (Fig. 5B: 2.01 million km^2^), when compared to present day (Figure 3B :1.99 million km^2^), but increased by 14.7% under the RCP8.5 SSP3 (Figure 5H: 2.29 million km^2^) scenario. Conversely, the total area where epidemics could start decreased under the RCP4.5 SSP1 by 47% (Figure 5C: 0.444 million km^2^), when compared to present day (Figure 3C: 0.836 million km^2^), but again increases under RCP6 SSP2 this time by 20.5%, and by 34% under the RCP8.5 SSP3 scenario (Figure 5F, Figure 5I).

**Fig. 5.**
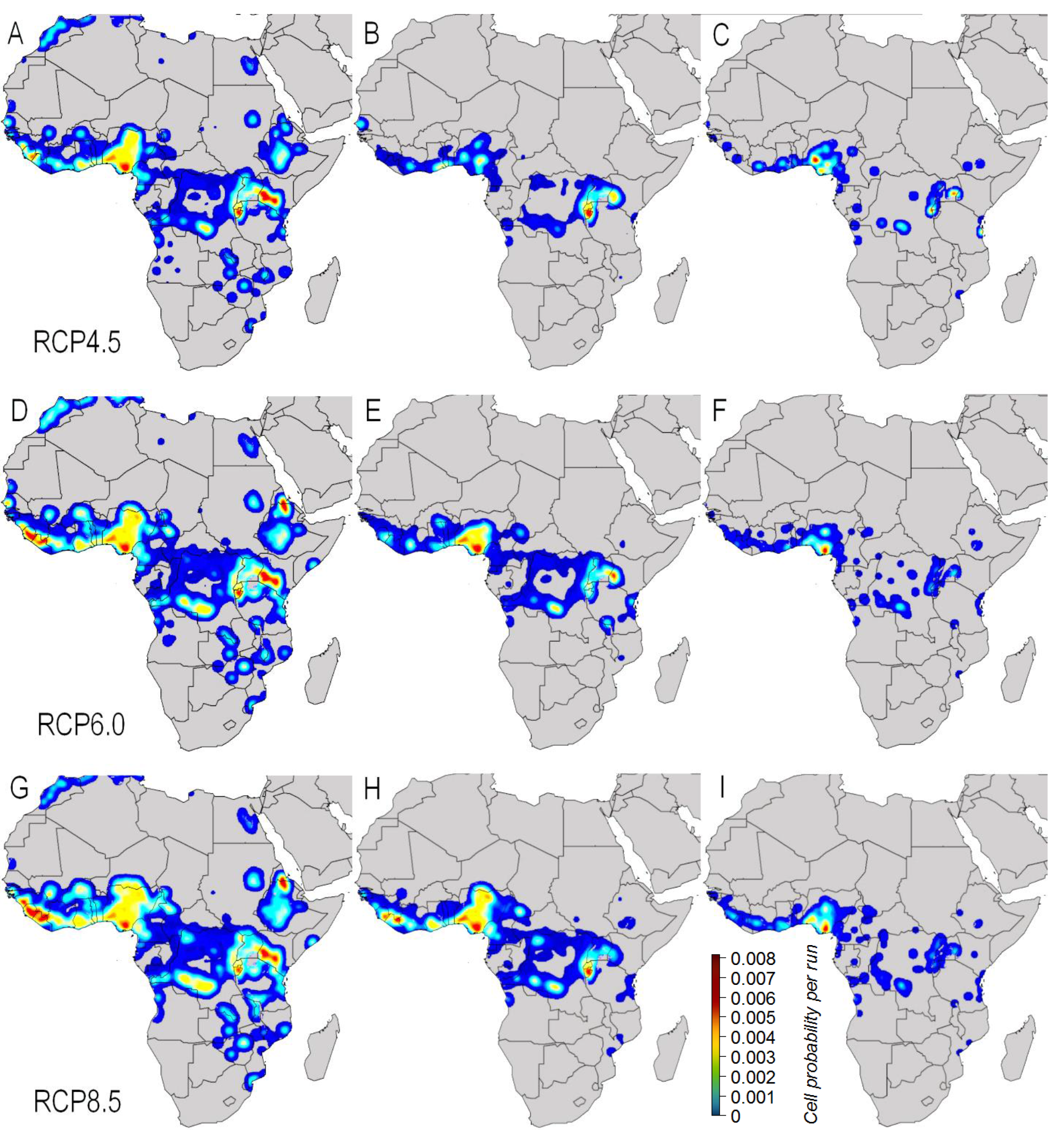
Future risk for Zaire Ebola virus (EBOV) from EMM simulations for 2070. Maps represent the proportion of times between zero (dark blue) and 0.01 (red) when a EVD-EBOV case was present in a grid cell (0.0416^°^), where (A, D, G) show all cases (both index and secondary), (B, E, H) index cases only, and (C, F, I) index cases from epidemics (1500+ cases), with data from EMM simulations for 2070, where rows show three different scenarios of global change (RCP4.5/SSP1, RCP6.0/SSP2, RCP8.5/SSP3).

The increases seen in the area affected is mirrored by greater total numbers of spill-overs experienced in future scenarios, with the greatest increase seen under the RCP8.5 SSP3 scenario at 7.92 (CI 7.62-8.19) spill-overs per year. Spill-over numbers increased with greenhouse gas concentrations (represented here by the RCP value) with a mean 0.257 spill over a year increase between the RCP4.5 SSP2 and RCP6 SSP2 scenarios, and a mean 0.343 spillover a year increase between the RCP6 SSP3 and RCP8.5 SSP3 scenarios. Greater increases were seen, however, with SSP change, with a mean 1.297 spill over a year increase between RCP4.5 SSP1 and RCP4.5 SSP2 and a mean 1.475 spill over a year increase between RCP6 SSP2 and RCP6-SSP3. In general, the probability of the index cases resulting in small outbreaks reduced in future environments, whereas the chance of epidemics increased (Figure 6). For instance, the proportion of epidemics per year (>1500 cases)decreased in the RCP4.5 SSP1 to 3.43% (from 5.8% in present day) but increased in all others, with RCP6 SSP3 gaining the greatest number, with epidemics in 9.5% of all simulations. The number of catastrophic epidemics (>2,000,000), generally increased with both RCP and SSP values up to 3.43% and 3.54% for the RCP6 and RCP8.5SSP3 scenarios respectively, but again saw a decrease from the present day level (2.3 %) to 1.19% for just the single ‘bestcase’ future scenario (RCP4.5 SSP1).

**Fig. 6.**
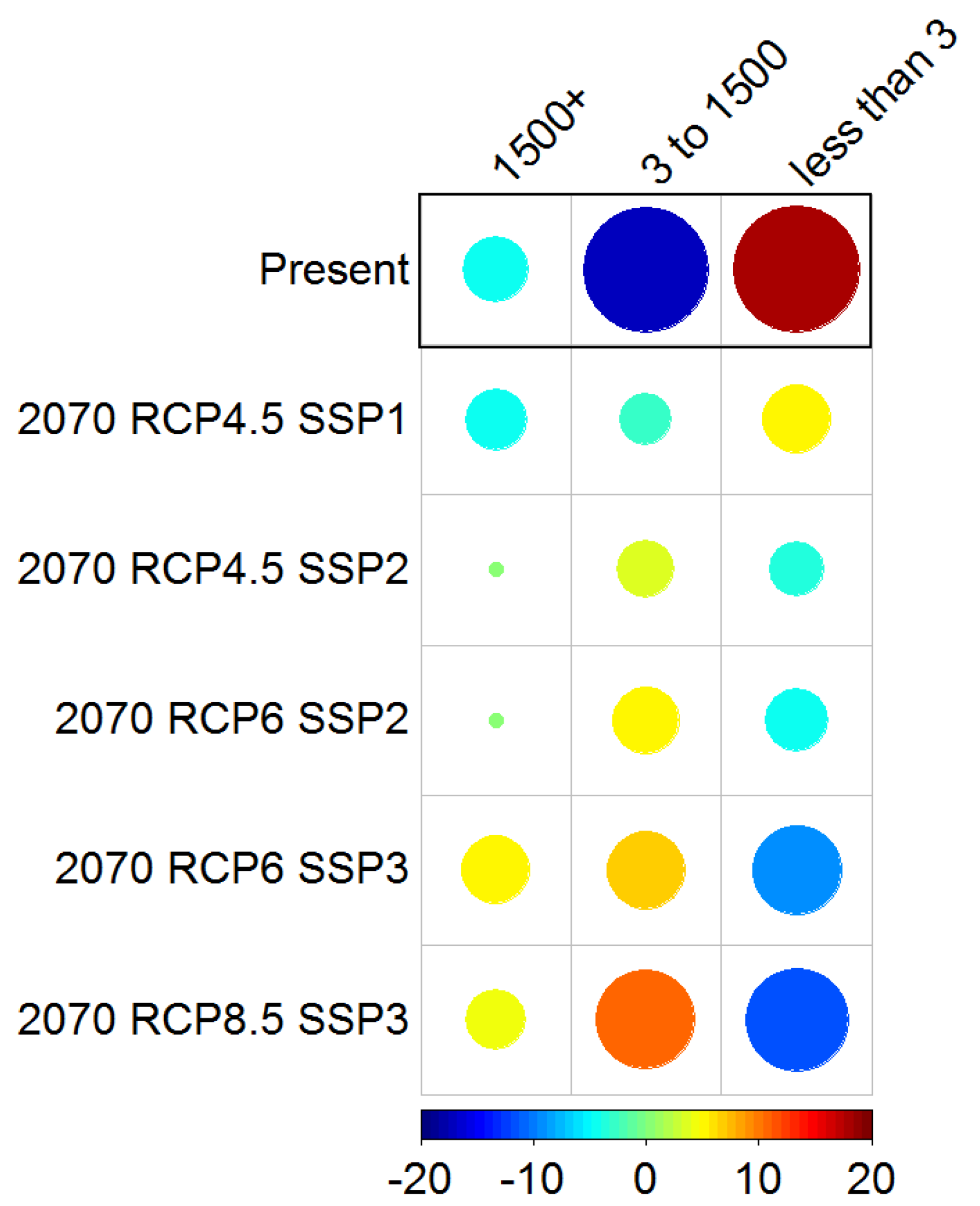
Comparison of 2070 EMM simulation scenarios by EVD-EBOV final epidemic size. Circles represents standardized residuals from a chi-squared test of association between simulation scenario and final outbreak size category. More orange/red colours show greater than expected number of outbreaks in a cell (for any given scenario and final outbreak size), with more blue colours representing fewer than expected outbreaks. Size of circle indicates the quantity greater or less than expected, with large circle more different than expected from random allocation of simulation runs among grid cells and small circles close tothe expected number.

## Discussion

We show that changes in future expected disease incidence are likely to be related to the rate of global environmental change. According to our study, EVD mitigation attempts would be best placed in efforts to reduce both population growth, increase socioeconomic development and ameliorate climate change, such that global change most closely tracks the RCP4.5 SSP1scenario. Global binding commitments to reducing climate change may act to slow the effects,but evidence (41) suggests a wholesale change is difficult. Expected decreases in poverty and a concomitant increase in healthcare resources, therefore, would appear to be the most realistic approaches to reduce the future EVD disease burden. While vaccinations may be effective, the sporadic nature of spill-over events mean it is unclear where vaccination should be targeted and whether it would be cost-effective at this time (35). More generally, increasing health care provision and poverty reduction efforts in West Africa would not only reduce the potential effects of EVD but also other diseases, including those that have yet to emerge in earnest, such as Marburg virus disease (42), Lassa fever (43), and Nipah/Hendra virus infection (44).This, in turn, could limit disease emergence to local outbreaks, preventing nosocomial infections and acting to prevent subsequent epidemics.

Changes to SSP scenarios, which control levels of poverty and human population size in our models, had a greater impact than changing the climate and land-use change (here mediated via RCP scenario). This is not surprising as poverty reduction increases the presumed EVD-EBOV healthcare response in our simulations, and many of the countries in the endemic region are expected to have substantial reductions in poverty levels by 2070 (37). Similarly, contact rates in our simulation (both between humans and between humans and wildlife) depend linearly on human population growth, whereas climate change increases EVD-EBOV cases through more complex interactions. Species distribution models indicate that the presumed wildlife hosts prefer warm and wet conditions (Figure S1-2), which are expected to increase in these regions according to the HADGem3-AO climate model (38)(Fig. S5). This expansion of the optimal conditions for presumed the wildlife host species effectively increases the at-risk human population by including more of the northern, eastern and southern areas of Africa (Figure 3A). Predicted future anthropogenic land-use changes, however, reduces the optimal wildlife host habitat, thereby reducing human-wildlife interactions.

We identify Nigeria as, not only a key area for epidemics to be initiated, but also an area with potential for many small outbreaks. This might indicate that our model has not correctly balanced the impact of healthcare infrastructure on disease spread, regional behavioural barriers to infection or regional differences in contact rates between both humans and hosts. Until these additional factors are explicitly tested, the high human density and known presence of putative wildlife hosts mean that this area should be consider at high risk of initiating epidemics.

There is a pressing need to better understand the spatial variation in other key disease transmission parameters. For instance, bush-meat hunting is an important process by which human populations come into contact with large bats resource (45) and the spatial variation in bush-meat extraction is likely a component of spill-over variation. Little is known, however, about bush-meat hunting outside a few specific studies but there is potential to use spatial interpolation techniques to make reasonable predictions in un-sampled areas. Our model does not incorporate this data or test its impact and, similarly due to lack of data resources, we do not use information about local differences in funeral practices. Hospital compartments are thought to be useful to understand quarantine and super-spreading events but there is very limited data on the quality and geographic reach of small health clinics. Some other important behavioural trends are not captured in our model, such as the post-outbreak behavioural reactions of human populations e.g. mass migration away from affected regions. Recent findings regarding the persistence of Ebola virus in semen of convalescent men may also help explain the intermittent spatiotemporal patterns of infections in endemic areas (46, 47). Future work incorporating such data, may further improve the spatial resolution and accuracy of risk estimates.

Our approach demonstrates not only an important framework understanding Ebola but also for other diseases. Analysing diseases singly cannot be an effective approach for policy making at a large geopolitical scale, particularly in regions with multi-disease burden and limited healthcare resources. Net disease risk patterns, when summed across a wide variety of zoonoses, will be an emergent property of the distribution of very different wildlife host species and their respective responses to increasing anthropogenic land-use conversion and climate change. Any lack of data in the short-term does not reduce the obvious importance of understanding future disease trends. Attempts, such as ours, establish a first heuristic step on a pathway to building intervention measures aimed at reducing overall future disease burden.

## Materials and Methods

### Environmental Mechanistic Model (EMM) EBOV

Using our discrete-time, stochastic epidemiological compartmental model incorporating spatial environmental variability (30), we extended the approach to not just simulate pathogen spill-over but also subsequent human-to-human transmission, focusing on the Zaire Ebola virus (EBOV) (Figure 2). Within grid cells (0.0416^°^) covering continental Africa, we used a Susceptible, Exposed, Infectious, Funeral and Removed (SEIFR) EVD-EBOV disease compartmental model (following13, 19, 23) to estimate the number of individuals per compartment, in each time step*t*, for present day bioclimatic, land use and demographic conditions. Although some previous compartmental models for EBOV have included a Hospital compartment (48), adding this complexity was not feasible over large and poorly known geographical areas. Without knowing more about the spatial variation in health seeking behaviour, exactly which grid cells contain clinics, and the variation of healthcare resources in these clinics, adding in this compartment would not likely significantly improve our model's ability to predict the progression of outbreaks. Further more, hospital interventions had the least impact controlling EVD outbreaks in a recent meta-analysis (24). All analyses were carried out in R v.3.2.2 (49). Each stage of the EMM simulation is discussed in more detail below:

#### Stage 1: SEIFR compartmental model within grid cells

We used starting EBOV transmission characteristics of incubation time = 7 days, onset of symptoms to resolution = 9.6 days, case fatality rate(CFR) σ = 0.78, and burial time = 2 days (23) to parameterize the SEIFR compartmental model to determine transition rates *α*(between Exposed to Infectious compartments), *γ*_*σ*_ (Infectious to Funeral),*γ*_*1−σ*_ (Infectious to Removed), and *γ*_F_(Funeral toRemoved) (Figure 2). Toincorporate sensitivity around these transmission parameters, we allowed values to vary for each simulation run by sampling from a Gaussian distribution where themean was their initialvalue and standard deviation was fifth of the mean, to give a reasonable spread of values. For each time step*t*, the number of individuals moving between all compartments was estimated by drawing randomly from a binomial distribution (Section S1 equation (1)), parameterized using the respective compartmental transition rates. Transition rates for compartments were assumed to be the same in all grid cells except for the transition between Susceptible to Exposed. The per grid cell Susceptible to Exposed transition rates were determined by the force of zoonotic infection *λ*_z_, and the force of infection *λ* (Figure 2) and these were calculated as follows:

##### (a) Force of Zoonotic Infection, *λ*_z_

The force of infection for zoonotic transmission *λ*_z_, per time step *t*, was estimated as the product of the probability of host presence *H*, and spill-over rate *κ* (Section S1 equation (2)). Without any evidence to the contrary (15, 50), we parameterized *H* by calculating the spatial probability of the presence of the most likely EBOV reservoir host species based on available data (Old World fruit bat species *Epomophorus gambianus gambianus, Epomops franqueti, Hypsignathus monstrosus,* and *Rousettus aegyptiacus* see Table S1) within each grid cell (0.0416°) across the African continent using species distribution models (SDMs) (51) and assuming constant pathogen prevalence. We also calculated the spatial probability of the presence of other species which are known to provide an alternative route of infection, but likely do not act as reservoirs (*Gorilla spp*., *Pan spp*., *and Cephalophus spp*.) (12). SDMs for each species were inferred using boosted regression trees (BRT) using distribution data from the Global Biodiversity Information Facility (GBIF) (52) and 11 presentday bioclimatic and landuse variables (Table S2). Data with coarse scale GBIF spatial coordinates (decimal degree coordinates with less than four decimal places)were filtered out of the analysis. To reduce spatial autocorrelationand duplicate records, any records that co-occurred in the same grid cell were removed. Lastly, GBIF records older than 1990 were discarded to ensure samples more closely matched the current landscapes. BRT tree complexity was set at5 reflecting the suggested value and the learning rate was adjusted until >1000 trees were selected (53). A total of 25 models were estimated for each species using four fifths of the distribution data asa training dataset andone fifth as a testing dataset, chosen randomly for each model. Thosewith the highest predictiveability (high area under operating curve, AUC and true-skill statistic, TSS values) were selected as the best model for each species (Figure S1). The most important spatial variables determining distributions across the different reservoir host species were BIO7 Temperature Annual Range, BIO13 Precipitation of Wettest Month, BIO2 Mean Diurnal Temperature Range and LandUse-Land Cover (Figure S2). The outputs from all putative reservoir (bat) species were combined into a single value representing the probability of any reservoir species being present and a similar approach was taken for the non-reservoir host species. The reservoir and non-reservoir host layers were then combined, but since onlya third of index cases were attributed to non-reservoir host spill-overs (10), wedown-weighted the probability of the non-reservoir occurrence by two thirds and reservoir occurrence by one third when combining the layers. The final resulting probability was bounded by zero and one. Additionally, as EBOV presence in non-reservoir host species is impossible without the presence of reservoir hosts, cells with a reservoir host probability of zero were given a value of zero irrespective of the non-reservoir host score. For computational simplicity, we assume that all human individuals have equal chance of exposure to infected host species. The initial value used for spill-over rate *κ*, per time step *t*, was estimated from the number of historic outbreaks *O* (defined here as distinct clusters of cases) (taken from empirical EBOV outbreak data 12), and the number of historically susceptible individuals *S*_h_ (inferred from human population estimates from 1976 to 2015 from 37) (see Section S1 equation (3)). During each simulation run, κ was allowed to vary using the same method as the compartmental transmission parameters above.

##### (b) Force of Infection, γ

The force of infection for human-to-human transmission *λ* per time step *t*, was estimated as the product of the effective contact rate *β*, and the number of individuals that can transmit the disease in each relevant compartment (*Infectious and Funeral*) per grid cell (0.0416°) (Section S1 equation (4)). We assumed that *β* for the Infectious and Funeral compartments was equivalent, due to the contact rates of moving individuals in the Infectious compartment being off set by large aggregations of individual sat funerals. We estimated the effective contact rate *β*, as the basic reproduction number *R_0_* divided by the product of the total number of individuals *N*, and infectious duration *D* (the sum of Infectious and Funeral compartment time, 11 days taken from 23). As a starting value for R_0_ we used avalue of 1.7 (54) and this was allowed to vary per simulation run using the same method as the compartmental transmission parameters above. As per previous research(30), we incorporated spatial variance in contact rates among grid cells usingaweighting factor *m*,where by the effective contact rate in grid cells with greater than expected contact rates was increased and decreased where fewer contacts were predicted(Section S1 equation (5)). We estimated *m* by creating an ideal free gas model of human movement within each grid cell and approximated collision frequency per person per day, using the following: the total individuals in each grid cell (estimated from Gridded Population of the World v3 55), an individual interaction sphere of radius 0.5 m, and using per person, daily walking distances in meters *vΔt*, where *v* is walking velocity, and*Δt* equals time period (Section S1 equation (6)). To capture geographic variation in human movement patterns, each grid cell was assigned a value for per person daily walking distance, based on the empirical relationship between daily walking distances and per person per country Gross Domestic Product (measured as Purchasing Power Parity from 37) (Table S3). As the availability of mass transit as alternative to walking tends to be centrally controlled, we assumed that grid cells in each country had the same value.

Under real conditions, the effective reproduction number *R*_e_decays over time as both efforts are made to control disease spread and as the pool of susceptible reduces, which results in *R_0_* being equal to *R*_e_ only when time step *t*is zero. Therefore, to calculate effective contact rate *β*, we allowed *R*_e_ to decay per time step *t* (Section S1 equation (7), equation (8) and equation (9)). However, countries that can invest more in health infrastructure (e.g., barrier nursing, surveillance) should see amore rapid reduction in *R*_e_ over time compared to countries that do not have such infrastructure and also a concomitantly, a decrease in CFR. Therefore we derived an empirical estimate of the relationship between wealth (measured using GDP-PPP per capita) and both the relative rate of decay of *R*_e_ over time (Section S1 equation 10) and CFR (Section S1 equation (11)), and using a spatially disaggregated poverty data layer (56) we weighted the per grid cell per time step *R*_e_ reduction and CFR accordingly to the values in each gridcell. While we found the relationship between wealth and both *R*_e_ and CFR reduction over time to be best described using curves with exponents of −0.08 and −0.02, respectively, this was inferred using relatively few data points (Table S4). In our simulation runs, therefore,we allowed these exponents to vary similarly to the parameters above, to allow either more linear declines or deeper curves to best estimate the true impact of this relationship.

#### Stage 2: SEIFR compartmental model between grid cells

We allowed those individuals that had contracted EBOV to travel between grid cells, specifically individuals in Exposed and Infectious (but not Funeral) compartments (Figure 2), but assumed for simplicity that the overall net movementof susceptible individuals between cellswas zero. As previously supported with empirical data, we employed a movement model that was weighted by both geographic distance and human density (31, 33) and was also geographically constrained to known transportation routes. The transmission rate *ϵ*, of individuals between target compartments of different grid cells was estimated by two different methods: between grid cells along road networks *ϵ*_r_, andalong flight routes *ϵ*_F_. We sampled randomly, froma binomial distribution, the number of travellers per grid cell and time step *t* (Section S1 equation (1)) with the probability of travel by road per day *ϵr*, being proportional to the distance to the nearest road using the Global Roads Open Access Data Set (Global Roads Open Access Data Set from 57). Global roads data set contains intotal 585413 routes from tracks to multi-lane highways and has been extensively validated for Africa (58). We allowed travellers to move freely (agnostic to any particular transportation method or country boundary) across the continent up to 10 road junctions in any direction from the centroid of the starting cell along the road network (Global Roads Open Access Data Set from 57), giving a potential of up to 500 km of linear travel per time step. Each proposed travel end point was given an individual probability from the daily distance travelled probability curve from (Figure 2(f) of 59), which is derived from transport data and validated against mobile phone data. For air travel, we set the potential pool of travellers as the individuals in grid cells containing airports across the world (from Open Flights Airport Database 60) plus all the Exposed individuals in the 8 grid cells surrounding each airport grid cell. We sampled randomly from a binomial distribution the number of travellers per grid celland time step*t* (SectionS1 equation (1)) with the probability of travel by air per day*ϵf*, being proportional to the total number of flights per day divided by the population of that country (37). We allowed travellers to move up to 2 edges on the current airline routes from airport origin using the (from Open Flights Airport Database 60). This approximates a traveller taking either a one or two-legged journey. Final destinations were sampled at random, based on all potential air routes having equal priority, but in most cases potential destinations were located nearby which by default meant that more distance travel was less likely than travel to a nearby location. For both road and air travellers, individuals were then added to the correct compartment of their final destination in the new grid cell and removed from the same compartment from the original source grid cell.

#### Stage 3: Impact of future anthropogenic change

##### (a) Future force of zoonotic infection*λ*_z_

We recalculated values of the force of zoonotic infection *λ*_z_, by estimating the probability of EBOV host presence, *H_2070_* under several different future integrated scenarios that incorporate projections of bioclimatic and land use variables (Table S2). Estimates of bioclimatic variables for 2070 were based on the HADGem3-AO climate model (61) under three Representative Concentration Pathways:RCP4.5, RCP6, and RCP8.5 (RCP45, RCP60 and RCP85 40). To estimate host presence probability in the future we needed to predict fine-scale future habitat data under theRCPscenarios. As only coarse categorisations are currently available (62), we therefore separately empirically estimated future land use-land cover (LULC) change (using MODIS data 36).Foreach grid cell we calculated the probability of each possible LULC change within the 2001-2012 MODIS dataset within a surrounding 5x5 cell grid using satellite data from 20. Based on these probabilities we simulated yearly LULC change across the region of interest for each grid cell from 2012 until 2070,and ran this simulation 100 times to create a bank of future possible landscapes, which were then summarized into three consensus landscapes representing low (with anthropogenic changes rejected where possible),medium by choosing the majority consensus across all 100 runs) and high anthropogenic change, (anthropogenic changes were chosen if available across the landscape) and we aligned these three scenarios to SSP1, SSP2 and SPP3 respectively.

##### (b) Future force of infection *λ*

Using predicted human demographicvariables and poverty levels for 2070, we recalculated values for the force of infection *λ*, by estimating the number of individuals per grid cell, *n* and effective reproduction number, *R*_e_. We inferred human population estimates per grid cell for 2070 by using the Gridded Population of the World v4(55) for present day and multiplying each cell by the expected future proportional change over that time period predicted by three Shared Socio-economic Pathways: SSP1, SSP2 and SSP3. Future poverty estimates per country were similarly inferred using a spatially-disaggregated GDP layer(63)multiplied by the expected change in per country GDP over the time period as predicted by the SSP integrated scenario. We note that as our travel probability is defined per person, increasing future populations will see aproportion increase in the amount of both road and air travel.

##### (c) Comparison of simulation runs

We reran the EMM simulations under 5 plausible combinations of 2070 future environmental-socioeconomic scenarios of global change and greenhouse gas concentrations: RCP4.5/SSP1, RCP4.5/SSP2, RCP6/SSP2, RCP6/SSP3, RCP8.5/SSP3 (64). These different input data options were,specifically: (i) RCP4.5-stabilization scenario in which total radiative forcing is stabilized shortly after 2100, (ii) RCP 6 - stabilization scenario in which total radiative forcing is stabilized short lyafter 2100, without overshoot, by the application of a range of technologies and strategies for reducing greenhouse gas emissions (iii) RCP 8 – worsening scenarios with increasing greenhouse gas emissions over time,leading to high greenhouse gas concentration levels, (iv) SSP1 – high regional cooperation, low population growth due high education and high GDP growth, (v) SSP2 – a ‘processes as usual’ scenario with ongoing levels of population growth and wealth,with medium estimates for both these by 2070, and (vi)SSP3 – regional antagonism, high population growth, unsustainable resource extraction and low economic growth. For each ofthe six scenarios we aimedfor 2500 runs of 365 days, each day measuring the number of spill-overs, the number of secondary cases associated with each spill-over, and the geographical areas affected. This allowed us to measure likelihood of spill-overs leading to small, medium and very large outbreaks, and also to determine the geographical areas with the highest risk of experiencing cases. We also noted the destination of any flights out of Africa that contained infected people.

## Acknowledgements

This work, Dynamic Drivers of Disease in Africa Consortium, NERC project no. NE-J001570-1was funded with support from the Ecosystem Services for Poverty Alleviation Programme (ESPA). The ESPA programme is funded by the Department for International Development (DFID), the Economic and Social Research Council (ESRC) and the Natural Environment Research Council (NERC). AAC is additionally supported by a Royal Society Wolfson Research Merit Award. We thank Prabu Sivasubramaniam for technical assistance, and A., Jones, M. Wilson, G. Mace, M. Leach,and C. Watts for comments on previous versions of the manuscript. All simulation data are available on figshare (figshare.com/s/c41c50a0675311e5b6b306ec4bbcf141) and the supplementary materials contain all other data. D.W.R. and K.E.J. developed the overall study design.D.W.R. carried out the modelling and data processing with assistance from K.E.J. All authors contributed to writing the manuscript. The authors declare no competing financial interests.

## Supplementary Information

### Section S1

#### EMM compartmental transition algorithms

For each time step t, the number of individuals moving through disease compartments both within and between grid cells (see Figure 2) was estimated using disease transmission parameters. We predicted the likely movement between disease compartments per time step, by drawing randomly from a binomial distribution. We describe this process below, using as an example the movement of individuals moving from Exposed to Infectious compartments within grid cells.

1. We determined the probability that a number of individuals were likely to move from the Exposed to Infectious compartments as:

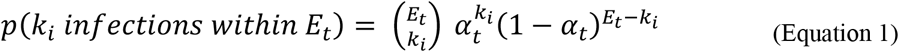

Where *k_i_* represents the number of individuals that enter theInfectious compartment, *^E_t_^* the number of Exposed individuals attime *t*, and *^α_t_^* the transition probability at time*t*.
2. Using equation (1), we determined for any value of *k_i_* the probability of *k_i_* individuals thatmove into the Infectious compartment, i.e., we computed the probability of the number of people, between 0 and the total number of individuals in the Exposed compartment,entering the Infectious compartment at time step *t*. We then drew randomly from this probability distribution to choose *k_i_* individuals that moved into the Infectious compartment, thereby weighting the choice towards the more likely outcomes given *α*.
3. Once the number of people that will be infected in the next time step ki was determined,then ki individuals were removed from the Exposed compartment and added to the Infectious compartment.
4. This process continued (per time step) until the number of individuals in the Exposed compartment equaled zero.

The same process was applied to every compartment change using the respective transition probabilities (i.e., substituting α in the above example). Movement of individuals between respective Exposed and Infectious compartments between grid cells was also modelled similarly, but stopping movements if the exposed or infectious number dropped to zero but with no change to susceptible numbers. Due to the high morbidity from this disease, individuals in the Infectious compartment were deemed less likely to travel and were awarded a travel probability that was half of the expected rate for non-symptomatic individuals.

#### Force of zoonotic infection*,^λ^*_z_ algorithms

The force of infection for zoonotic host to human transmission, *λz* was estimated per grid cell, per time step *t,* as follows:

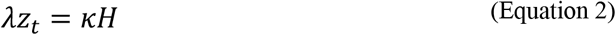

where *Κ* = spill-over risk, and *H* = probability of zoonotic host presence per grid cell. Spill-over event probability, Κ per person, per time step is given by:

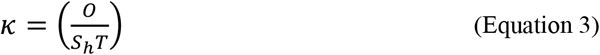

where *O* = number of historic outbreaks, *^S_h =_^*number of historically susceptible individuals, and *T* = total time when infections could have occurred. Note: Above we are estimating the probability of an individual being involved in a spill-over event directly from an animal host, which is distinct from the overall risk of contracting the disease.

#### Force of infection, *λ* algorithms

The force of infection for human-to-human transmission, per grid cell and per time step *t,* was estimated as:

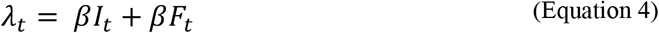

Where *β* = effective contact rate, *I_t =_*number of individuals in Infectious compartment at time step*t,*and *Ft* = number of individuals in Funeral compartment at time step *t*. For simplicity we assumed that *βI* and *βF* were the same (hereafter referred to as *β)*. When *t* = 0, *β*is given by:s

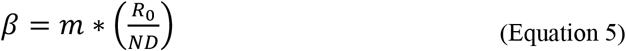

Where *R*0 = basic reproduction number, *m* = mobility, *N* = population size per time step, and *D* = duration in days that an individual is infectious. In this context, *m* was used to modify the ideal free gas model of human movement with distances travelled which are spatially variable across the landscape. We calculated a two-dimensional collision frequency *c*, per person per grid cell(65) as follows:

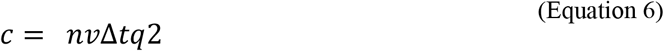

where *n* = number of individuals, *v* = walking velocity, *Δt* = time period and *q* = interaction sphere radius. In the context of our simulation, *v*Δ*t* represents daily walking distance. Then we defined *m* as the inverse deviation from a mean of *c* such that areas with more movement have a higher effective contact rate. However, when *t >*0 we redefined *β* as follows:

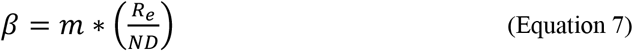

where *^R_e =_^*effective reproduction number, *m* = mobility, *N* = population size per time step, and *D* = duration in days that an individual is infectious. *R*_e_ is related to *R*0 but due to changes in human behaviour and health care responses, *^R_e_^* may be lower and decline over time, in addition to the implicit reduction in *R* as the pool of susceptibles decreases during an outbreak. We make the assumption that the effective reproduction number reduces on a daily basis due to increasingly strong health care responses over time.

So initially, when *t* = 1:

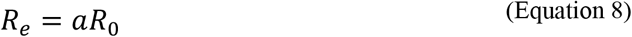

where *Re* = effective reproduction number at *t* = 1, *a* = decay rate, and *R*0 = basic reproduction number. However, when *t*>1:

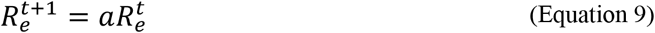

where 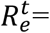effective reproduction number at time *t*, and *a* = decay rate. We define decay rate *a* per grid cell, from the empirical relationship between wealth and health outcomes. Using either direct or derived empirical estimates of the gradient of the change in *R*_e_ over time from (13, 19, 21, 22), we fitted an exponential decay curve between estimatesof per captia Gross Domestic Product measured as Purchasing Power Parity (from 37) and the gradient of *^R_e_^*change per day. The starting *Re* decay value *a* per grid cell, was given by:

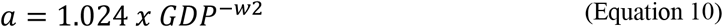

where the best estimate for exponent w2 was −0.848, *GDP* = Gross Domestic Product from (63), pseudo *r*^2^ = 0.76, and *n* = The poverty-weighted Case Fatality Rate (*w*CFR) per grid cell, was given by:

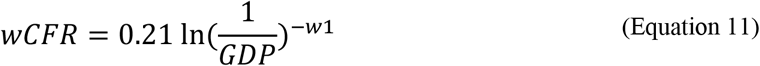

where the best estimate for exponent w1 was -0.0239, GDP = Gross Domestic Product from (63), pseudo *_r^2^_* = 0.9081, and *n* = 20.

**Fig. S1.**
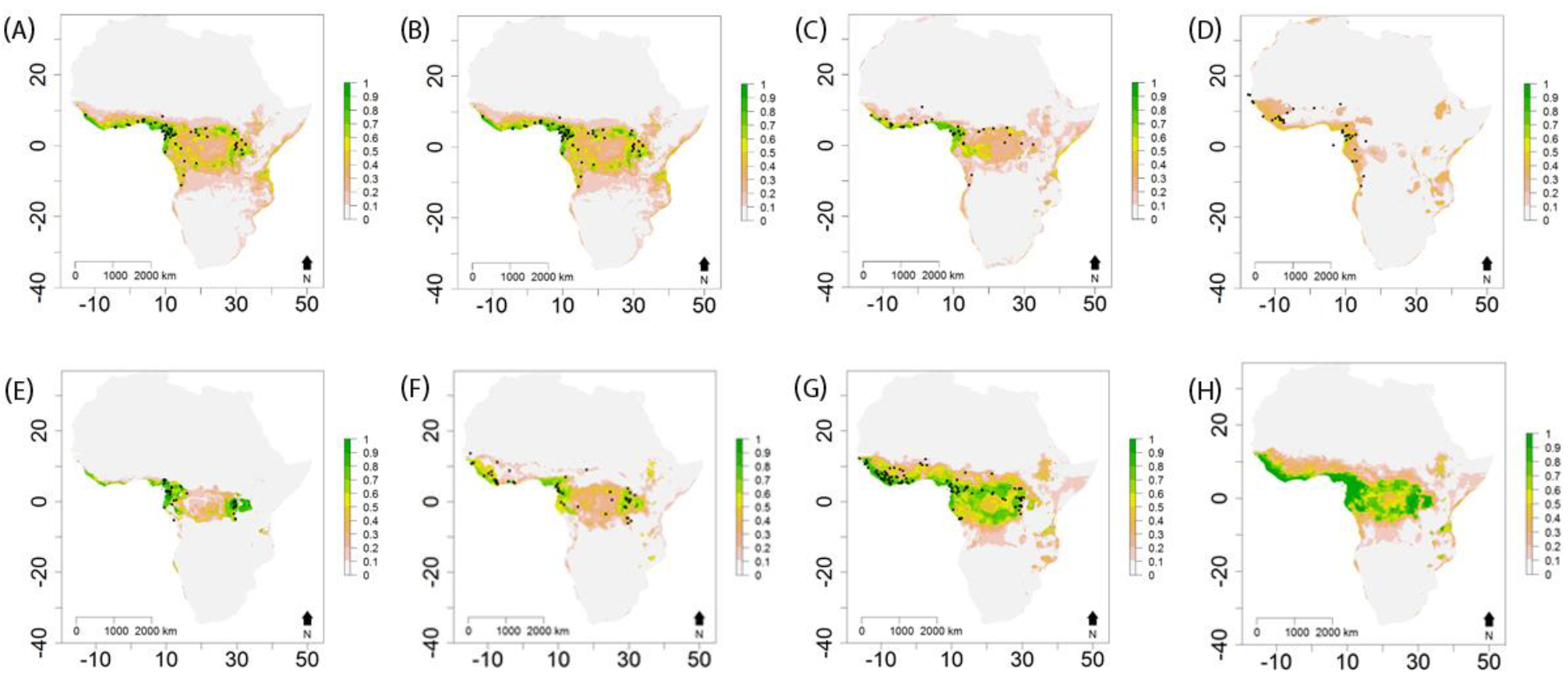
Maps of present day occurrence probability, H of EBOV host and other infection source species estimated from boosted-regression trees (BRT) models. Probability of species occurrence per grid cell (0.0416¯) is represented on a linear color scale where green is most suitable (p(H) = 1) and white unsuitable (p(H) = 0) where (A) Epomophorus gambianus gambianus; (B) Epomops franqueti; (C) Hypsignathus monstrosus; (D)Rousettus aegyptiacus; (E) Gorilla spp.; (F) Pan spp.; (G) Cephalophus spp.; and (H) allspecies combined. Axislabels indicate degrees in a World Geodetic System 84 projection. Filled black circles represent GBIF (52) occurrence records.

**Fig. S2.**
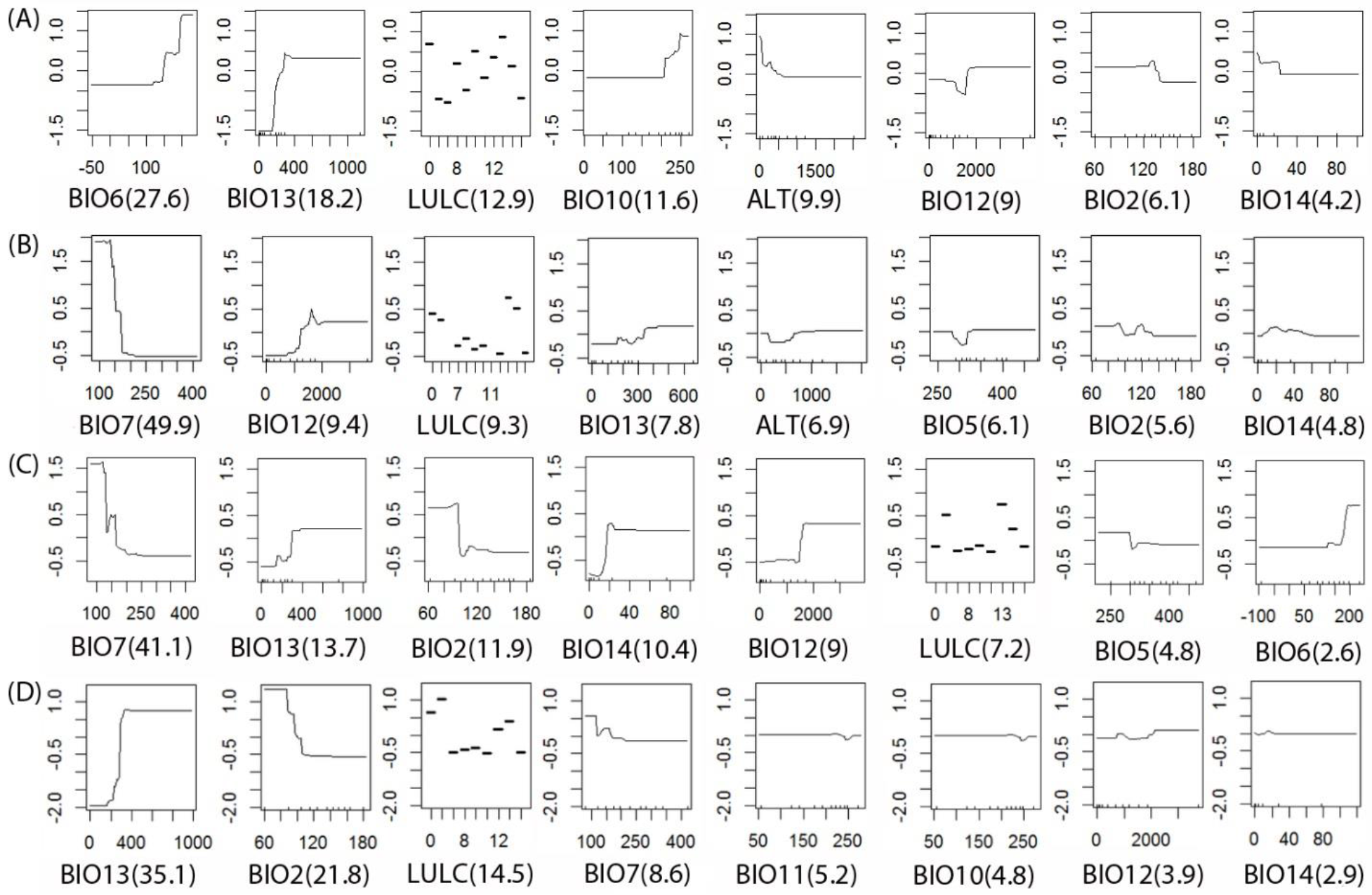
Response curves from boosted-regression trees (BRT) models of EBOV host species occurrences. Each plot represents the shape of the normalized fitted functions for each variable where (A) Epomophorus gambianus gambianus; (B) Epomops franqueti; (C) Hypsignathus monstrosus; and (D) Rousettus aegyptiacus. The relative percentage contribution of each variable to the model in terms of variance explained is given in parenthesis, where only the top eight variables of the model are included for each species. Variable abbreviations are defined in table S2.

**Fig. S3.**
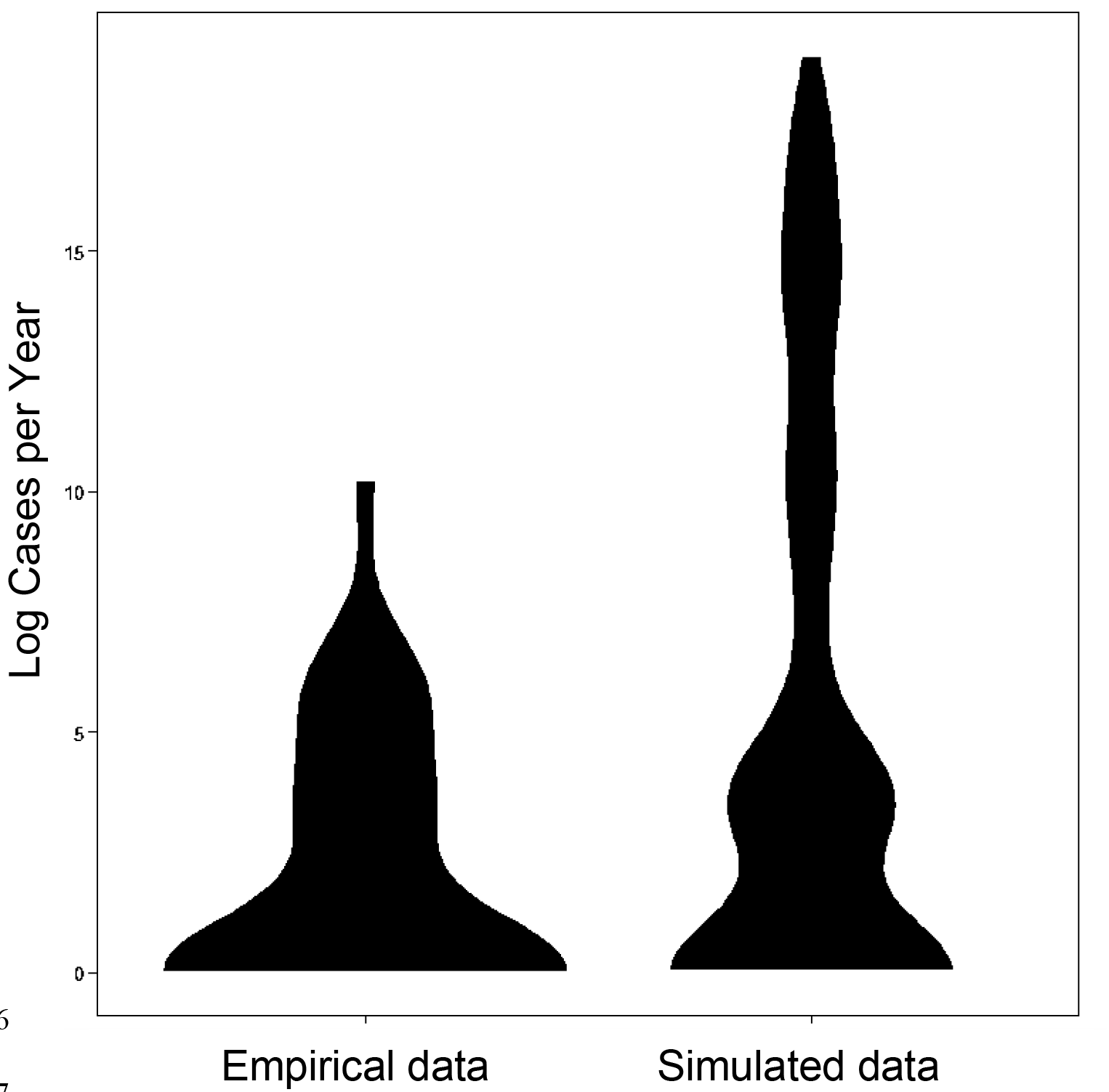
Distributions of relative frequency of EBOV cases per year. Violin plots represent the empirical observed (n=23 outbreaks) data of log total number of cases per year from 1967-2016 (66), and log total number of cases per year (n=2500 runs) from EMM simulations for present day environmental and demographic conditions.

**Fig. S4.**
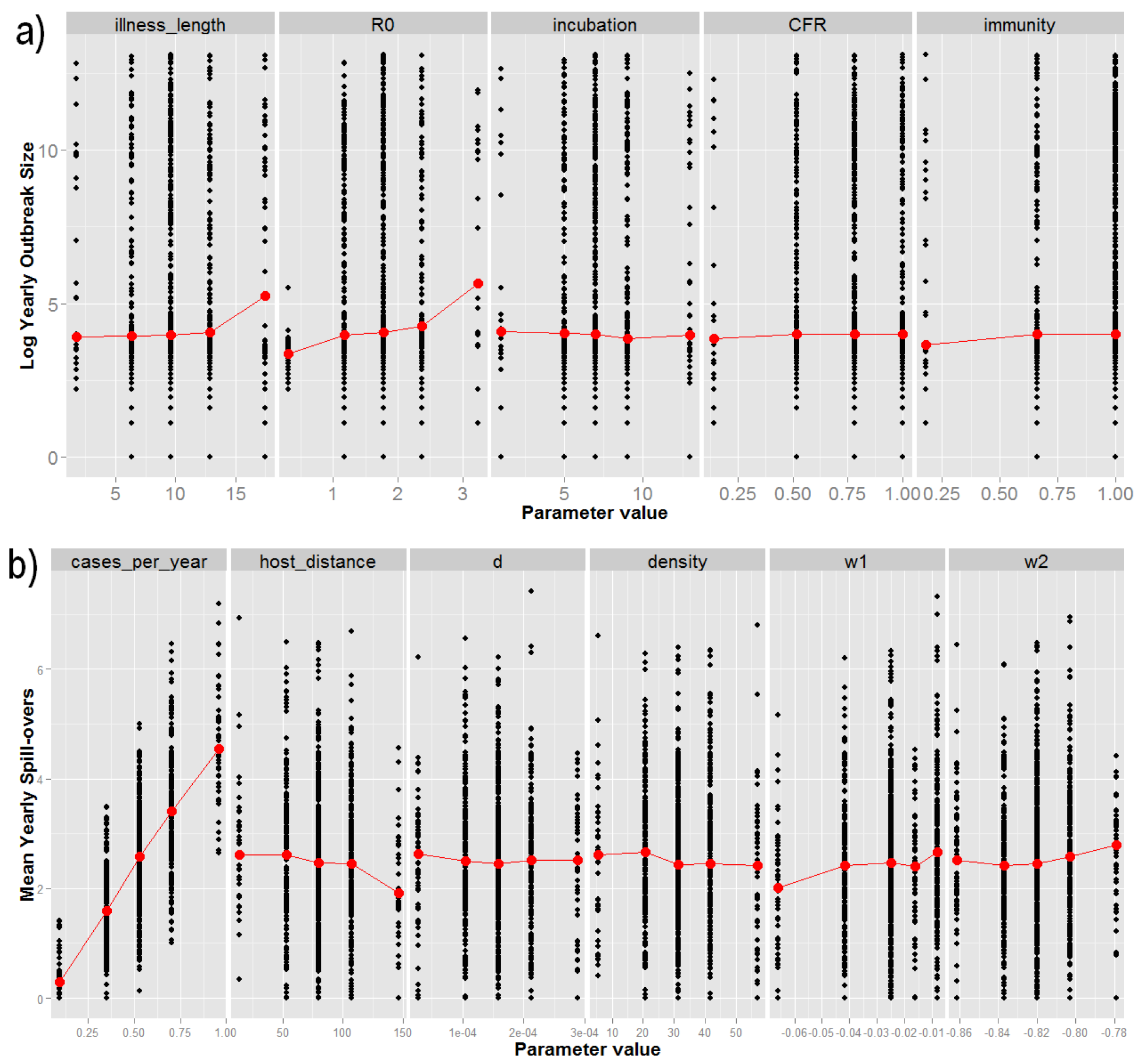
Sensitivity plots of input parameters for (a) total number of annual log EVD cases, and (b) mean annual spill-overs. Black dots show the response values per simulation and are jittered for greater clarity. Red dots represent the median values for each parameter value, and red lines join the medians to aid interpretation of any trend. Parameters are as follows: illness length - mean number of days in the infectious compartment; ^R_0_^ basic reproductive number; incubation - mean number of days in the exposed compartment; CFR - mean case fatality rate per illness; immunity - mean immunity to re-infection where 1 is totally immune; cases per year - mean spill-over rate constant; host distance - mean daily distance (m) travelled by host reservoir species; density - mean numberof reservoir host individuals per grid cell; w2-shape of the effective reproductive number (^R_e_^) decay curve, where low values represent a less curved, more linear negative relationship; and w1 - shape of the CFR~poverty curve, where lower values represent a less curved, more linear negative relationship.

**Fig. S5.**
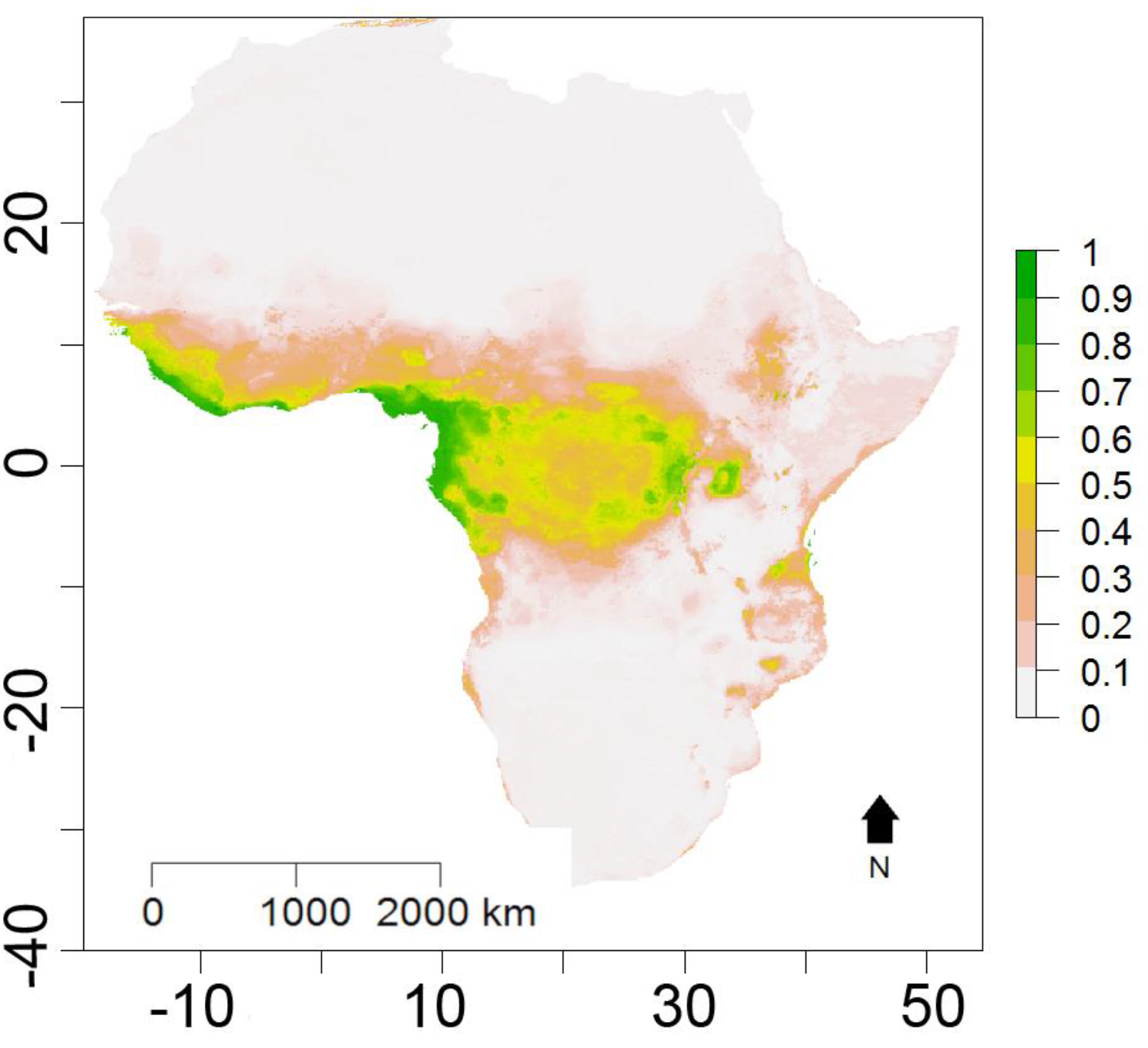
Map of future occurrence probability, H2070 of EBOV host and other infection source species estimated from boosted-regression trees (BRT) models under the medium outlook RCP6 scenario. Probability of species occurrence per grid cell (_0.0416^°^_) is represented on a linear color scale where green is most suitable (p(H) = 1) and white unsuitable (p(H) = 0) for all species combined. Axis labels indicate degrees in a World Geodetic System 84 projection.

**Table S1.**
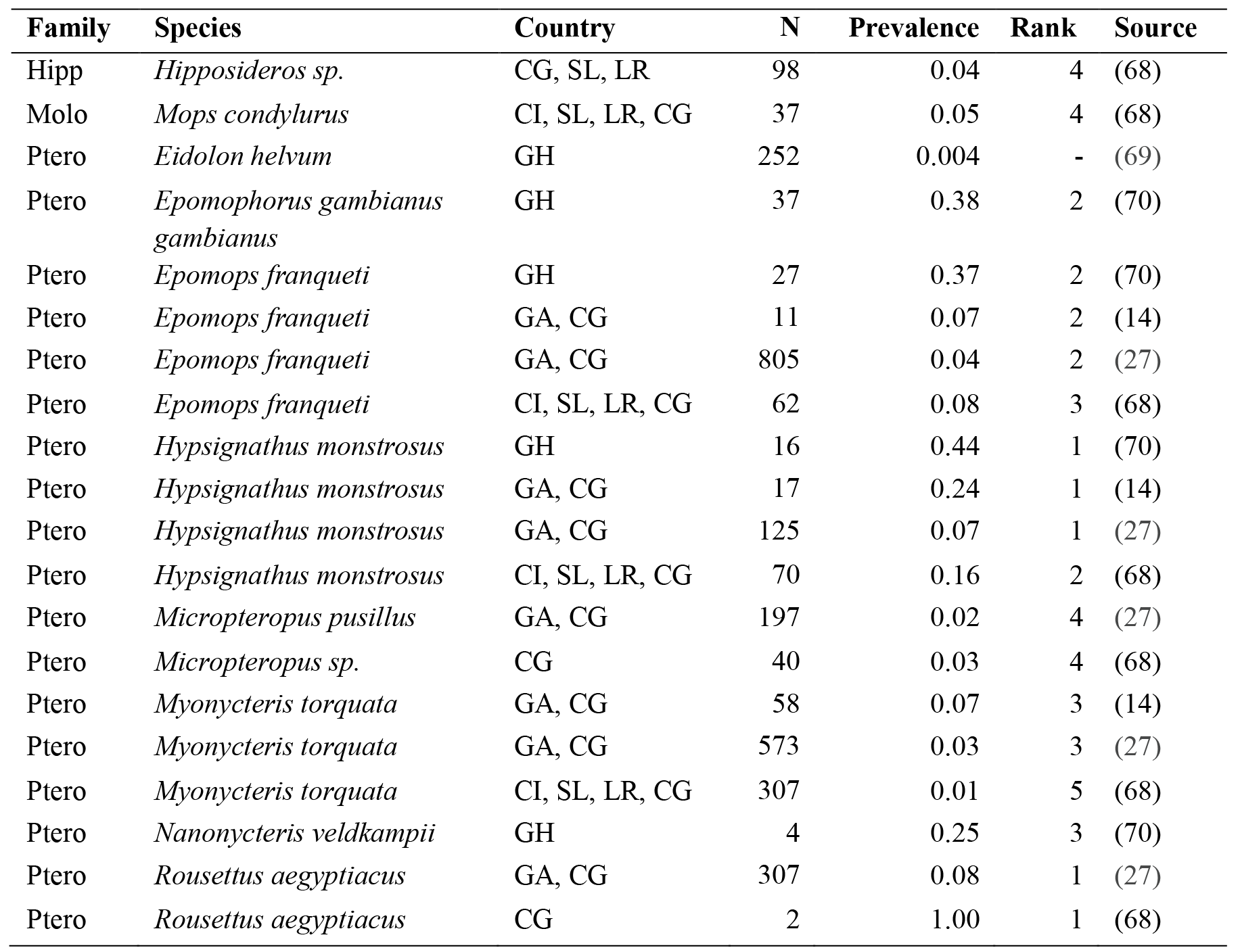
Seroprevalance of EBOV in reservoir host species. Species assignments followed the taxonomy in (67). Prevalence was measured as the proportion of positive results per sample and raw prevalence data was transformed to a rank within each study. Direct prevalence comparisons were not possible due to methodological differences. We estimated the most important EBOV host species as those that appear as the top two ranks in all sources. We identified four candidate bat species hosts: *Epomops franqueti, Epomophorus gambianus gambianus, Hypsignathus monstrosus, and Rousettus aegyptiacus*. N represents sample size; Hipp Hipposideridae; Molo Molossidae; Ptero Pteropodidae; CI CÔte d'Ivoire; SL Sierra Leone; LR Liberia; GH Ghana; CG Congo; and GA Gabon.

**Table S2.** Details of bioclimatic and land use variables used to estimate probability of EBOV host presence, H. Nine most orthogonal (<75% correlation) bioclimatic variables were chosen from (71). For analysis, all variables were reduced in latitudinal extent to 85° N, 58° S and resampled to a 0.0416° grid cell size using a World Geodetic System 84 projection. LULC is a categorical dataset where the most predominant land use-land cover type in each grid cell is given within the following categories: Evergreen needle leaf forest; Evergreen broadleaf forest; Deciduous needle leaf forest; Deciduous broadleaf forest; Mixed forest; Closed shrublands; Open shrublands; Woody savannah; Grassland; Permanent wetlands; Cropland; Urban and built-up; Cropland/natural vegetation mosaic; Snow and ice; Barren or sparsely vegetated; and Water bodies.

| No. | Variable Description | Original Spatial Extent | Original Spatial Resolution (cell size at equator) | Temporal Resolution | Source |
| --- | --- | --- | --- | --- | --- |
| 1 | BIO2 Mean Diurnal Temperature Range | Global | 1km | 2012 | (71) |
| 2 | BIO5 Maximum Temperature of Warmest Month | Global | 1km | 2012 | (71) |
| 3 | BIO6 Minimum Temperature of Coldest Month | Global | 1km | 2012 | (71) |
| 4 | BIO7 Temperature Annual Range | Global | 1km | 2012 | (71) |
| 5 | BIO10 Mean Temperature of Warmest Quarter | Global | 1km | 2012 | (71) |
| 6 | BIO11 Mean Temperature of Coldest Quarter | Global | 1km | 2012 | (71) |
| 7 | BIO12 Annual Precipitation | Global | 1km | 2012 | (71) |
| 8 | BIO13 Precipitation of Wettest Month | Global | 1km | 2012 | (71) |
| 9 | BIO14 Precipitation of Driest Month | Global | 1km | 2012 | (71) |
| 10 | ALT Digital Elevation Model | Global | 1km | 2008 | (72) |
| 11 | LULC Land Use-Land Cover | Global | 500m | 2001-2012 | (36) |

**Table S3.** Estimates of global daily walking distances, vΔt. Estimatesof daily walking distances were collected from the literature per country. Daily step numbers were converted to distance (km) using an average step length of 1.41m (73). As studies have suggested that daily walking distance is stratified among income categories (74), countries were assigned to income bands based on per capita Gross Domestic Product (GDP) (measured as Purchasing Power Parity from 37) such that the poorest countries were given a value of 1 and the richest 4.A mean estimate of walking distance was calculated for each band. Countries were then assigned a walking distance corresponding to their GDP band. No estimates were found forband 3 ($1600 - $35000), so countries in this band were given daily walking distances halfway between bands 2 and 4.

| Country | Steps | Distance (km) | GDP band | GDP PPP Per capita (lower bound) \$ | GDP PPP Per capita (upper bound) \$ | Mean km Per GDP Band | Source |
| --- | --- | --- | --- | --- | --- | --- | --- |
| Niger | - | 7 | 1 | 0 | 1600 | 9.6 | (75) |
| Central African Republic | - | 8 | 1 | 0 | 1600 | 9.6 | (75) |
| Chad | - | 15 | 1 | 0 | 1600 | 9.6 | (75) |
| Mali | - | 13.2 | 1 | 0 | 1600 | 9.6 | (75) |
| Niger | - | 4.8 | 1 | 0 | 1600 | 9.6 | (75) |
| South Africa | 12471 | 8.85 | 2 | 1600 | 13000 | 8.5 | (76) |
| Tanzania | - | 8.3 | 2 | 1600 | 13000 | 8.5 | (77) |
| Australia | 9695 | 6.88 | 4 | 35000 | 128530 | 5.6 | (78) |
| Japan | 7168 | 5.08 | 4 | 35000 | 128530 | 5.6 | (78) |
| Switzerland | 9650 | 6.85 | 4 | 35000 | 128530 | 5.6 | (78) |
| United States | 5117 | 3.63 | 4 | 35000 | 128530 | 5.6 | (78) |

**Table S4.** Collated epidemiological data on EBOV outbreaks. Data on 19 locations that have experienced EBOV outbreaks or importations and have data on either Case Fatality Rate (CFR) (13, 18, 54, 79–83) or on Effective Reproductive Number change (21, 54, 79, 84–86) (^R_e_^ gradient per week). The latter data was either taken directly from tables or text from within literature sources or estimated (Spain, United Kingdom, Nigeria, United States) from descriptions of outbreak events detailed in the sources. Child mortality data for the year of outbreak is taken from WorldBank Development Indicators (37)

| Location | County | Year | ln GDP per capita<br>for year | CFR | $R_e$ gradient per week |
| --- | --- | --- | --- | --- | --- |
| United States | Texas | 2014 | 4.74 | 0.3 | 0.5 |
| Guinea |  | 2014 | 3.09 | 0.707 | 0.113636 |
| Sierra Leone |  | 2014 | 3.31 | 0.69 | 0.076923 |
| Liberia |  | 2014 | 2.99 | 0.723 | 0.04 |
| Germany |  | 2014 | 4.66 | 0 |  |
| Spain | Madrid | 2014 | 4.53 | 0 | 0.5 |
| United Kingdom | London | 2014 | 4.59 | 0 | 3 |
| Nigeria |  | 2014 | 3.77 | 0.666667 | 0.533333 |
| Mali |  | 2014 | 3.24 | 0.75 |  |
| Congo, Dem. Rep. |  | 1976 | 2.72 | 0.88 | 0.105 |
| Gabon |  | 1994 | 4.14 | 0.61 |  |
| Congo, Dem. Rep. |  | 1995 | 2.72 | 0.81 |  |
| Gabon |  | Early-1996 | 4.17 | 0.68 |  |
| Gabon |  | Late-1996 | 4.17 | 0.75 |  |
| Gabon |  | 2001–2002 | 4.15 | 0.82 |  |
| Congo, Rep. |  | 2001–2002 | 3.58 | 0.76 |  |
| Congo, Rep. |  | Early-2003 | 3.6 | 0.89 |  |
| Congo, Rep. |  | Late-2003 | 3.6 | 0.83 |  |
| Congo, Rep. |  | 2005 | 3.65 | 0.75 |  |

